# Mobbing-like response to secondary predator cues is not a form of teaching in meerkats

**DOI:** 10.1101/2020.07.02.182394

**Authors:** Isabel Driscoll, Marta Manser, Alex Thornton

## Abstract

Across many taxa, individuals learn how to detect, recognise and respond to predators via social learning. Learning to recognise and interpret predator cues is essential in the accurate assessment of risk. Cues can come directly from a predator’s presence (visual, acoustic) or from secondary predator cues (SPCs, such as hair/feathers, urine or faeces) left in the environment. Animals show various responses to encountering SPCs, which are thought to act in reducing risk to the individual. Meerkats, Suricata suricatta, show a response to SPCs not described in any other species: they display a mobbing-like behaviour. The function of this behaviour is unclear as unlike mobbing, the response it so closely resembles, it cannot serve to drive predators away. We used experiments to investigate whether adults may use this mobbing-like response to teach naïve young how to recognise and respond to predators. Meerkats are known to teach pups hunting skills, but there is as yet no evidence that any species other than humans teaches across multiple contexts. We used experimental presentations of SPCs to test whether wild adult meerkats respond more intensely to SPCs in the presence of naïve pups, as would be expected if the behaviour serves to promote learning. Contrary to this prediction, response intensity was lower when pups were present than when they were absent, and declined as the number of pups in the group increased, possibly due to costs associated with foraging with dependent young. Response intensity instead increased with increasing group size, number of group members interacting with the cue, and varied with predator cue type. These results suggest that the mobbing-like response to SPCs is not a form of teaching in meerkats. Instead, this behaviour may function to increase the recruitment of others to investigate the SPC. Exposing group members to SPCs may better inform them of the nature of the threat, facilitating more effective defensive group responses.

## Introduction

The ability of prey animals to mount appropriate defensive behaviours in the face of predation is vital to survival. Accurately assessing current predation risk aids in informing risk-appropriate behaviours, limiting unnecessary time and energy expenditure on non-acute or non-immediate threats. Individuals can gauge predation risk through personal assessment of the current situation and from the risk assessments of others, by using social information (Dall *et al.* 2005; Crane & Ferrari 2013). Access to social information is thought to be a key benefit of group living, aiding in detecting, recognising and responding appropriately to predators. In animals across many taxa, social learning plays an important role in shaping the development of appropriate responses to predators (see reviews: Griffin 2004; Crane & Ferrari 2013). One common antipredator behavioural response that is often learnt via social learning is mobbing (Curio *et al.* 1978a; Davies & Welbergen 2009; Cornell *et al.* 2012; Feeney & Langmore 2013; Griesser & Suzuki 2017).

Mobbing is a method of predator deterrence which involves individuals gathering around and investigating a potential threat, and in many species is accompanied by the production of distinctive calls (Curio *et al.* 1978b; Graw & Manser 2007). Mobbing is conspicuous and costly in terms of time and energy expenditure, advertises an individual’s location, and may increase the risk of injury or death (Curio *et al.* 1978b; Krama & Krams 2005; Tórrez *et al.* 2012), but it can also provide important advantages. For instance, mobbing may offer opportunities for individuals to learn to recognise and respond appropriately towards predators by observing conspecifics’ behaviour. Naïve juvenile Siberian jays, *Perisoreus infaustus*, for example, learnt to both recognise and mob a predatory goshawk, *Accipiter gentilis*, following a single observation of a knowledgeable individual mobbing the predator (Griesser & Suzuki 2017). However, the principal benefit of mobbing is thought to be predator deterrence, either by intimidating and driving away the predator, or by alerting it that it has been detected and thus reducing the chance of successful attack (Abolins-Abols & Ketterson, 2017; Caro, 2005). While the benefits of mobbing and driving a predator away are clear, meerkats, *Suricata suricatta,* also exhibit a rather perplexing form of behaviour, where they show mobbing-like responses towards secondary predator cues (SPCs).

Secondary predator cues are cues left in the environment by predators; such as fur, urine, faeces, feathers, scent markings and regurgitation pellets, sometimes referred to as either direct or indirect cues (Persons *et al.* 2001; Severud *et al.* 2011; Nersesian *et al.* 2012; Zöttl *et al.* 2013). These cues can indicate predator presence in the vicinity and provide information about the nature of the threat. In most cases prey avoid SPCs or respond with defensive behaviours such as increased vigilance (Monclús *et al.* 2005; Zidar & Løvlie 2012; Garvey *et al.* 2016; Tanis *et al.* 2018), reduced activity (Persons *et al.* 2001; Sullivan *et al.* 2002; Lehtiniemi 2005), refuge use (McGregor *et al.* 2002; Sullivan *et al.* 2002; Ferrari *et al.* 2006; Belton *et al.* 2007), and moving away from the cue (Amo *et al.* 2004; Shrader *et al.* 2008; Mella *et al.* 2014). However, some species respond by approaching and inspecting SPCs, presumably to gain further information about the source of the cue (Belton *et al.* 2007; Furrer & Manser 2009; Zöttl *et al.* 2013; Garvey *et al.* 2016; Collier *et al.* 2017). Some species are able to ascertain the type of predator (Van Buskirk 2001; McGregor *et al.* 2002; Mella *et al.* 2014), predator size (Kusch *et al.* 2004), age of the cue (Barnes *et al.* 2002; Zöttl *et al.* 2013; Kuijper *et al.* 2014) and the predator’s diet from these cues (Mathis & Smith 1993; Apfelbach *et al.* 2015). Meerkats take this inspection behaviour one step further by responding to SPCs in a very similar manner to that shown when they mob real predators. To our knowledge, meerkats are the only species to show such mobbing-like responses to SPCs. Other mongoose species, such as dwarf and banded mongooses, do recruit to and inspect SPCs (Furrer & Manser 2009; Collier *et al.* 2017), however, meerkats show a more overt, higher arousal, behavioural response. When meerkats encounter SPCs they approach and investigate the cues, raising their tails, piloerecting (raising their fur) and making recruitment calls. These responses are all characteristic features of meerkat mobbing behaviour (Graw & Manser 2007), but in contrast to true mobbing, the mobbing-like response towards SPCs serves no function in deterring predators (see figure 1 for comparison). The potential benefit of responding to a SPC as if it were the predator itself is thus very unclear, particularly given that the response is conspicuous and involves time and energy costs. One potential function of the mobbing-like response towards SPCs by meerkats could be to act as a form of teaching for naïve young.

**Figure 1.**
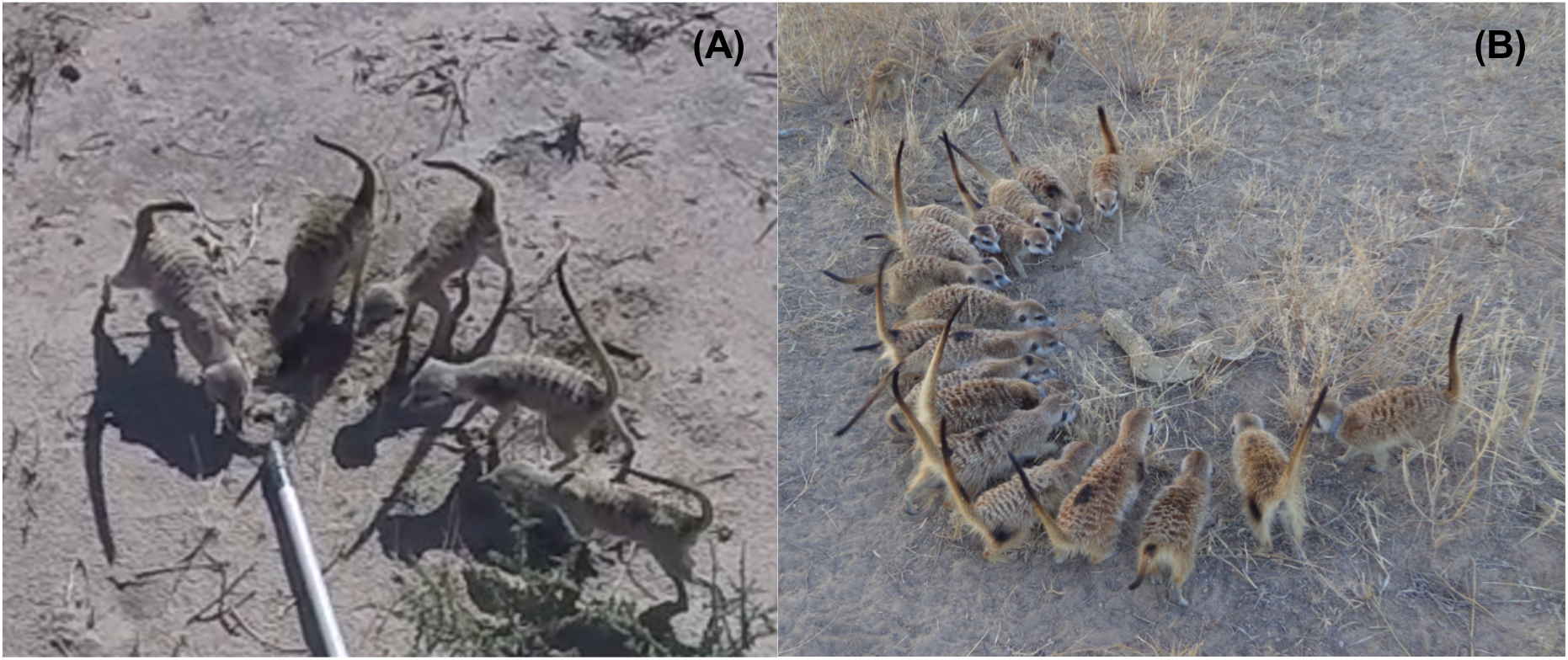
(A) Meerkats’ mobbing-like response to experimental SPC presentation, (B) meerkats’ mobbing response to a predatory puff adder, Bitis arietans, (Photo: Jess Snow)

Teaching is a form of active social learning whereby knowledgeable individuals invest in promoting learning by the naïve (Thornton & Raihani 2008). According to established operational criteria, teaching involves (i) an individual, A, modifying its behaviour in the presence of a naïve observer, B, (ii) A incurs a cost or no immediate benefit by doing so, (iii) as a result of A’s behaviour B acquires a skill or knowledge faster than it would have otherwise, if at all (Caro & Hauser 1992). Teaching was once regarded as uniquely human, but there is now strong experimental evidence for teaching in a handful of non-human animals including meerkats (Thornton & McAuliffe 2006), tandem-running ants, *Temnothorax albipennis* (Franks & Richardson 2006) and some species of birds (Raihani & Ridley 2008; Kleindorfer *et al.* 2014; Chen *et al.* 2016).

In stark contrast to human teaching, all known cases of teaching in other species occur in a single context. Meerkats, for example are known to teach pups to handle difficult prey items by gradually introducing them to live prey (Thornton & McAuliffe 2006), but there is no evidence of teaching in other contexts (Thornton 2008; Thornton & Malapert 2009). Thus, if the mobbing-like response to SPCs serves in part as a form of teaching, then this would provide the first evidence for teaching in multiple contexts outside of humans. Passive social learning may be sufficient to learn about SPCs through group recruitment events. However, the unusual mobbing-like response in meerkats raises the possibility that there is an additional aspect of this behaviour. Specifically, by inspecting and responding conspicuously to SPCs in the presence of pups, adults could incite naïve pups to approach investigate the cue themselves. Exaggerated mobbing-like responses could therefore provide valuable opportunities for pups to learn about predator characteristics (e.g. odour) and appropriate behavioural responses in a relatively safer environment.

In this study we used experimental presentations to investigate whether the mobbing-like response to SPCs functions as a form of teaching in wild meerkats. Meerkats are cooperatively breeding mongooses from the arid regions of southern Africa, in which all group members help to rear dependent pups (aged 0-3 months) (for more detailed information see: Clutton-Brock & Manser 2016). Meerkat pups make extensive use of social information in the development of foraging skills and anti-predator responses (Hollén & Manser 2006; Hollén *et al.* 2008; Thornton & Clutton-brock 2011) and are known to learn hunting skills via teaching (Thornton & McAuliffe 2006). We tested whether mobbing-like responses towards SPCs may constitute another form of teaching in animals, with adults modifying their behaviour so as to promote learning in pups. Specifically we predicted that, as per the first criterion of Caro and Hauser’s definition of teaching (Caro & Hauser 1992), adults should increase the intensity of their mobbing-like response (raised tails; piloerection; recruitment calls; (Graw & Manser 2007)) would be greater when pups were present and when cues were novel to the pups but not to adults.

## Methods

### Study site & species

Experiments were carried out on six groups of wild meerkats at the Kalahari Meerkat Project in and around the Kuruman River Reserve, South Africa (Clutton-Brock *et al.* 1998). All members of the population used in the experiments were habituated to observations at < 1m, with individuals identifiable from unique dye marks on their backs (Jordan *et al.* 2007). Group sizes ranged from 3-24 and the life history of all group members were known as part of long-term study of the population for over 20 years.

### Cues

We presented two different cue types: (1) domestic cat, *Felis catus,* urine samples, obtained from local veterinary surgeries during medical procedures and stored in the freezer and (2) African wildcat, *Felis lybica*, fur samples, obtained from a recently deceased individual found (within 6 hours of death) on the reserve and stored in the freezer. Both domestic cats and wildcats are common predators on the reserve. Adults were likely to have encountered the predators and their associated cues previously but, given the frequency of predator encounters, it was highly likely that pups were naïve. Pilot studies determined that adults responded to both predator cues with a mobbing-like response. Samples were portioned into 5mls of urine and 0.1g of fur and stored at −20°C. To ensure that meerkats were responding specifically to the cues and not the experimental set-up, equivalent quantities of water and dry grass were used as matched controls for the urine and fur respectively. We removed cues from the freezer to defrost 2-3 hours before presentation, keeping them in a cool bag with ice blocks until presentation and wore latex gloves to avoid contaminating the cues with human scent.

### Presentations

We conducted presentations while the group were foraging in the morning. The first trial at a group was after pups had been born, but were still being babysat at the burrow, and had not begun foraging with the group (no pups: NP). This allowed conditions to be kept as similar as possible across trials (including hormonal changes associated with reproductive events), while still allowing comparison of trials with and without pups. Pups began foraging with the group at around three to four weeks of age, but initially spent much of their time in sheltered locations (e.g. in boltholes or under bushes) begging for food and did not participate in group alarm or mobbing events. The second trial, with pups present (pups present 1: PP1) was conducted when pups were approximately six-seven weeks (21 ± 3 days after they began foraging with the group) and spent the majority of the time actively moving between helpers. Subsequent trials (pups present 2 and 3: PP2 and PP3) were conducted at one week (7 ± 1 day) intervals. For trials 1-3 (NP, PP1, PP2) the same cue type was used and for trial 4 (PP3) a different cue was used, representing a novel cue (Table 1). We predicted that adults would show the lowest mobbing intensity to PP2 as the cue type was not novel to pups or adults. Half of the groups were presented one combination of cues (Group A – urine, urine, urine, fur) and the other were presented the opposite (Group B – fur, fur, fur, urine). For each trial a cue was presented and a control, with a randomised order of predator or control presentation. The second cue was presented ten minutes after the interaction with the initial presentation had ended.

**Table 1.** Set up of the four experimental trials showing the conditions, cue type, cue novelty, pup age and pup location.

|  | Trial 1 – NP |  | Trial 2 – PP1 |  | Trial 3 – PP2 |  | Trial 4 – PP3 |  |
| --- | --- | --- | --- | --- | --- | --- | --- | --- |
| **Treatment** | No pups |  | Pups present 1 |  | Pups present 2 |  | Pups present 3 |  |
| **Cue** | A. Urine | B. Fur | A. Urine | B. Fur | A. Urine | B. Fur | A. Fur | B. Urine |
| **Cue novelty?** | N/A |  | Yes – to pups |  | No |  | Yes – to pups |  |
| **Pup age** | 24 days ± 3 days | | 49 days ± 3 days | | 56 days ± 3 days | | 63 days ± 3 days | |
| **Pups foraging?** | Pups babysat at burrow |  | Foraging with group for 21 days |  | Foraging with group for 28 days |  | Foraging with group for 35 days |  |

Cues were presented 30 minutes after the group had left the burrow in the morning to begin foraging, and after at least 10 minutes of normal foraging behaviour following an alarm event, so as to minimise the effect of any previous stress on responses to the presentation. The cues were presented in a petri dish filled with sand at the end of a 1 m pole, to reduce association of cues with the human presenter. One week prior to beginning the experimental presentations the cue presentation apparatus was presented to all group members filled only with sand to habituate them to the set up and ensure that responses during the experimental trials were to the cue and not the apparatus. At the start of each trial, we presented the relevant cue to a randomly selected target individual (non-pup, > 6 months) from the group. If the individual did not initially respond to the cue, we presented it again up to three times. If this still did not elicit a response the cue was presented to another randomly chosen individual to prevent over-exposing any one individual to the cue. A trial began once an individual responded to and began interacting with the cue. Trials were conducted at least one week apart to reduce possible habituation to the cues. Presentations were video recorded using a GoPro (Hero 4) and audio recorded holding a microphone (Sennheiser ME 66 with a K6 powering module, sampling frequency 44.2 kHz, 16 bits accuracy) connected to a recorder (Marantz Solid State Recorder PMD661 MKII) at a distance of approximately 1-1.5m from the cue presentation (*see supplementary material for video example*).

### Behavioural analysis

Video recordings were coded using the open-source software BORIS (Friard & Gamba 2016), noting the behaviours of each individual that interacted with the cue. Details and definitions of the behaviours recorded are given in Table 2. Only the behaviours of individuals that interacted with the cues were recorded. Presentations that elicited no response from the initial target individual were not included in the analysis unless subsequent presentations to the rest of the group did not elicit a response.

**Table 2.** Ethogram of the behaviours analysed in response to the secondary predator cue presentations.

| Behaviour | Description |
| --- | --- |
| **Interact** | *Duration of time spent interacting with the cue, when the individual was within 0.3 m of the cue following initial approach and exhibiting one of the following behaviours (indicating a direct interaction): facing the cue directly, touching and sniffing the cue, rocking back and forth facing the cue, tail raised, and piloerecting. Interaction ended when an individual was quadrupedally vigilant (scanning on four legs), on bipedal vigilance (scanning on two legs), or resumed foraging. A new interaction began if the individual started interacting again.* |
| **Tail raise** | *Tail raised vertically above the body within 0.5 m of the cue (within close proximity). Recorded the duration of time until the tail was lowered below horizontal with the body.* |
| **Piloerection** | *Fur visibly raised within 0.5m of the cue (within close proximity). Recorded the duration of time until the fur was no longer visibly raised.* |
| **Recruitment Call** | *The recruitment call type (low or high urgency) given in response to the cue presentation.* |

### Acoustic analysis

Acoustic recordings were analysed using RavenLite to determine the type of recruitment call given (high or low urgency) in response to the presentation (Bioacoustics Research Program 2016). We recorded the duration of calling bouts and classified the urgency of recruitment calls based on the acoustic structure (outlined and defined in: Manser 2001; Manser *et al.* 2001). Due to the nature of the audio recording method it was not possible to determine which individual was calling or how many individuals were calling, as typically several individuals were recruited to the cue simultaneously.

### Statistical analysis

Statistical analysis was conducted with R version 1.1.463 (R Core Team 2015), using the packages *lme4* for Generalised Linear Mixed Model (GLMM) analyses. Group identity was fitted as a random term for analysis of group-level responses (analysis *a*), and individual and group ID were fitted as random terms in analyses of individual responses (*b-g*). Analyses were conducted on the behavioural responses of all non-pup (group members > 3 months; hereby referred to as adults) individuals present for the experimental predator cue presentations. Model assumptions were checked using residual plot distribution techniques. We applied an information theoretic (IT) approach for model selection, using Akaike’s information criterion (AIC) to rank the models following the approach used by Richards *et al.* (Richards *et al.* 2011). Models within AIC ≤ 6 of the model with the lowest AIC value formed the ‘top set’. We then applied the ‘nesting rule’ to the top set, removing more complex versions of nested models from the top set so as to not retain overly complex models.

All models (*a-g*) included the explanatory terms: treatment (NP, PP1, PP2, PP3), cue type (fur, urine), number of pups (0-3 months old), and number of adult (> 3 months) group members. As individuals’ responses may have been influenced by their group mates’ behaviour, we also fitted the proportion of the group interacting with the cue (*b-g*) and the highest urgency level of call type heard in the group before each individual was recruited as additional explanatory terms (*a-g*). As the original target individual to whom the cue was presented could not, by definition, have heard any prior calls made in response to the cue, call type was categorised as target individual, no call, low urgency or high urgency. Individual age, sex, and dominance rank were initially included in the models but removed to reduce model complexity, as they never ranked in the top five models with the lowest AIC values during model selection. *A priori* combinations of fixed effects were used in model building based on biological-relevance.

As the number of pups in the NP treatment was, by definition, zero, the effects of treatment and number of pups could be correlated. To address this, we also ran the analysis with the results of the NP treatment excluded. The results of these models were qualitatively very similar to those conducted on the full dataset (see supplementary material, Table 1).

### Group-level response

First, we analysed the influence of treatment, cue type, number of non-pup group members and number of pups on the group level response (*model a*). We used a GLMM with binomial error structure and logit link function, fitting the proportion of the number of individuals responding as the numerator and the total number of other adults present in the group as the denominator, to take into account variation in group size. For this analysis we grouped the recruitment events with low urgency calls and no calls given, to allow model convergence as there were only two instances of recruitment following no recruitment calls. These two categories were grouped as they were both representative of a lower perceived risk.

### Individual response

We then used GLMMs to examine the factors influencing individual behaviour. We conducted a GLMM with binomial error structure and logit link function to examine how the explanatory terms outlined above, influence whether or not an individual interacted with the cue using a binary (0/1) response term (*model b*). We excluded the response of the original target individual presented to from this analysis as this interaction signified the beginning of the trial. Among those individuals that did interact, we examined the factors influencing the duration of interactions using a GLMM with a gamma error structure, for over-dispersed continuous data (Zuur *et al.* 2009; Richards *et al.* 2011), and log link function (*model c*). We also examined whether or not each of these interacting individuals raised their tail as a binary response term (0/1) using a GLMM with binomial error structure and logit link function (*model d*). For model d we grouped low urgency and no recruitment calls to allow model convergence, as there were only two instances of individuals raising their tails following no recruitment calls. Among those individuals that did raise their tails, we examined the factors influencing the duration of individual’s tail raising using a GLMM with a gamma error structure and inverse link function (*model e*). We also examined whether or not the interacting individuals piloerected as a binary response term (0/1), using a GLMM with a binomial error structure and logit link function (*model f*). This analysis did not include call type, as no individual showed piloerection if no recruitment calls or low urgency calls had been heard in the group. Among those individuals that did piloerect, we examined the duration of piloerection using a GLMM with a gamma error structure and log link function (*model g*).

### Responses of pups

At least one pup interacted with the cue presentation in 14/18 trials. Of 51 observations, representative of every pup in every trial contributing an observation, there were 19 instances of pups interacting with the predator cues. On average 1.06±0.78 (range: 0 to 3) pups were recruited to the predator cues. Pups’ interactions lasted an average of 46.10±9.02 seconds. Among the pups that did interact 15/19 raised their tails for on average 24.40±8.51 seconds, and 5/19 piloerected for on average 14.36 ±4.78 seconds.

### Responses of adults to control vs experimental stimuli

In response to experimental SPCs individuals typically displayed a combination of responses of: approaching the stimuli, investigation of the cue (visually assessing, touching with paws and sniffing), recruitment calling, tail raising, piloerection, and in some cases head bobbing and rocking body movements. In total there were 48 cue presentations analysed (combined predator and control). For six out of the 24 predator cue presentations analysed, cues need<u>ed</u> to be presented more than once to elicit a response. There was one instance in which all group members did not respond following three SPC presentations to each member of the group, the trial to the original target individual was used for the analysis. In one case the original target individual did not respond to the cue, but another individual came and investigated the cue independently and recruited other group members, this trial was also included in the analysis. Individuals never reacted to control presentations with more than a brief investigation and only those directly being presented with the control ever interacted with it. No recruitment calls were given to control cues and no individuals were recruited. Of the 24 control presentations 19 initial target individuals interacted with the control cue, as defined in Table 2, and five did not interact with the cue at all after being presented to three times. Of those that did interact with the control cue, interactions lasted on average 3.77±0.63 seconds, ranging between 0.75-11.25 seconds. Of the 19 individuals that did interact with the control cue only 4 raised their tails for an average of 3.88±1.16 seconds and none piloerected. Mean interaction duration with predator cues (29.66±2.64 seconds), ranging between 1.75-131 seconds, lasted approximately eight times longer than control cue interactions (paired t-test, t_23_ = 6.587, p < 0.001). Control presentations were not included in the models due to this consistent lack of response.

### Group-level responses to SPCs

#### (a) Proportion of the group recruited

On average a proportion of 0.34±0.02 of all non-pup group members were recruited to the predator cue presentations following the response of the initial interacting individual, and this depended on the number of pups present in the group. GLMM analyses produced six models in the top set, of which one (model a.5; supplementary material 1 Table 3) was retained with the lowest AIC value. This model contained only the number of pups present in the group as a negative predictor of the proportion of the non-pup group members recruited (GLMM: estimate (SE) = −0.201(0.107), *χ*^2^= 3.810, p = 0.05; Fig.1; supplementary material 1 Table 2). Call type appeared in the second highest-ranked model, but did not have a robust effect (GLMM: estimate (SE) = 0.567 (0.573), *χ*^2^= 1.260, p = 0.26).

**Figure 1.**
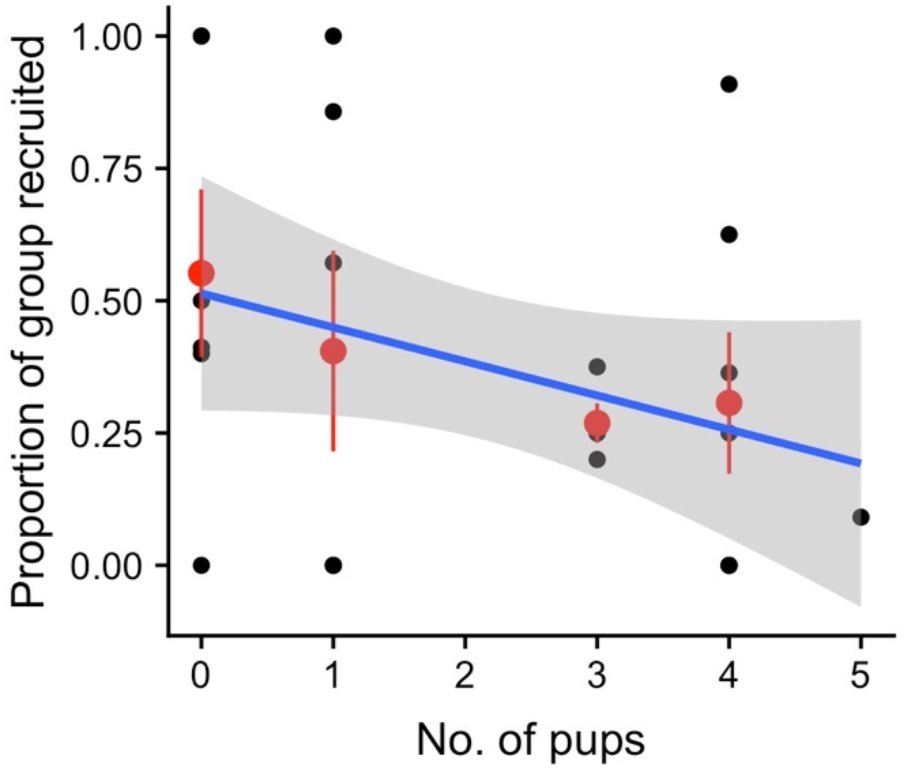
The overall proportion of the non-pup group members recruited dependent on the number of pups present in the group (n = 24 presentations). Red points indicate the mean proportion recruited with error bars signifying standard error. Blue logistic regression line with the shaded area illustrating the 95% confidence interval.

### Individual responses to SPCs

#### (b) Interacted (y/n)

Of the 202 observations, representative of every individual in every trial contributing an observation, 92 individuals interacted with the predator cue. Out of these 92 cases, 22 were the original target individuals to whom the cue was presented and the remaining 70 were subsequent recruits. GLMM analyses produced three models in the top set, of which one (model b.10; supplementary material Table 4) was retained following the application of the nesting rule. This model contained only the proportion of the group already interacting with the cue as a positive predictor of whether each new recruit interacted with the cue itself (GLMM: estimate (SE) = 2.992 (0.817), *χ*^2^= 14.753, p < 0.001; Fig.2A; supplementary material Table 2). Call type and treatment (models 9 and 11; supplementary material Table 5) appeared in the second and third highest-ranked models respectively, but neither factor appeared to have a robust effect (GLMM: Call type: *χ*^2^= 1.906, p = 0.39; Treatment: *χ*^2^= 2.732, p = 0.43; supplementary material Table 5).

**Figure 2.**
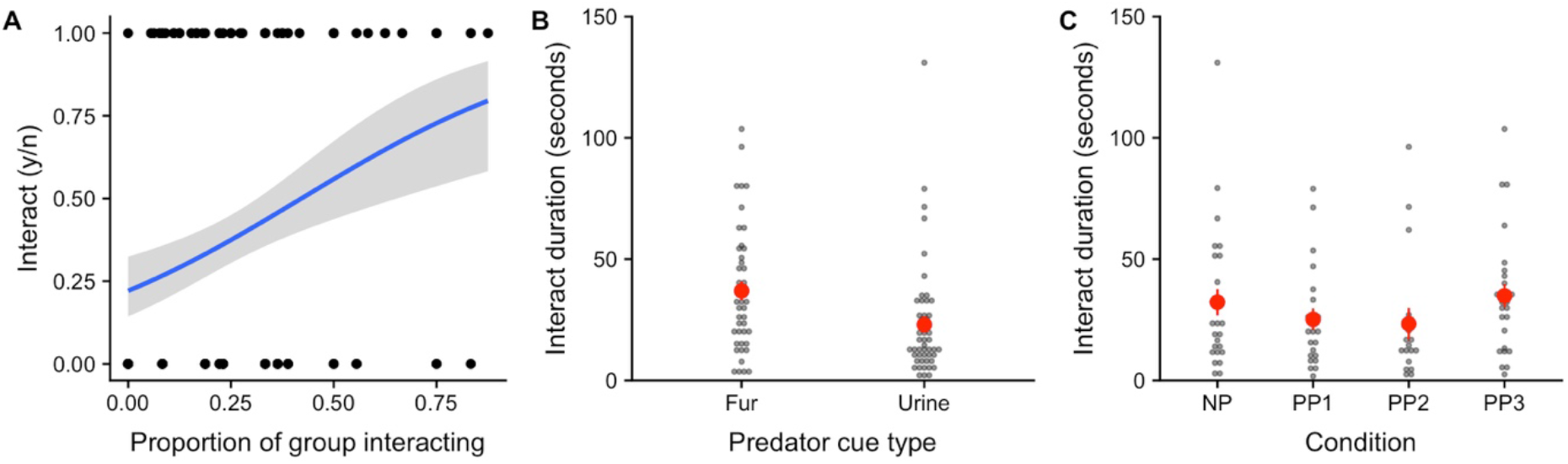
(A) The likelihood of an individual interacting with the cue yes (n = 92) or no (n = 110) dependent on the proportion of individuals in the group already interacting with the cue presentation prior to an individual beginning their interaction. Blue logistic regression line with the shaded area illustrating the 95% confidence interval. (B) The interaction duration in seconds of individuals that interacted with the presentation cues for the two cue types, fur (n = 44) and urine (n = 48), and (C) for each condition (no pups (n = 27), pups present 1 (n = 22), pups present 2 (n = 17), pups present 3 (n = 26)). Red dots indicate the mean interaction duration for each cue type with error bars signifying the standard error.

#### (c) Interaction duration

Individuals interacted with the predator cues for on average 29.66±2.64 seconds. GLMM analyses produced three models in the top set, of which one (model 5; supplementary material Table 6) was retained following the application of the nesting rule. This model contained only the predator cue type presented, with individuals interacting longer with fur cues, 36.92±3.81 seconds, than urine cues, 23.00±3.40 seconds (GLMM: estimate (SE) = −0.511 (0.169), *χ*^2^= 8.787, p = 0.003; Fig.3B; supplementary material Table 2). Treatment appeared in both the second and third highest-ranking models; when included with number of pups present, with both factors appeared to have an important effect (model 3; treatment: *χ*^2^= 10.89, p = 0.01; number of pups: estimate (SE) = 0.243 (0.107), *χ*^2^= 5.156, p = 0.02; Fig.3C supplementary material Table 6). However, when treatment was included with cue type, the effect of treatment was not robust (model 6; treatment: *χ*^2^= 4.979, p = 0.17; Fig.3C; supplementary material table 7). Interaction durations were greatest in NP (32.25±5.44 seconds) and PP3 (34.90±4.87 seconds), when cues were novel to the group, and lower in PP1 (25.18±4.50 seconds) and PP2 (23.32±6.54 seconds) when cues were not novel. NP differed most from PP2 (effect size = 0.35, t = −2.19, p = 0.03; supplementary material Table 7), and less from PP1 (effect size = 0.28, t = −1.09, p = 0.27; supplementary material Table 7) and PP3 (effect size = −0.10, t = −0.63, p = 0.53; supplementary material Table 7).

**Figure 3.**
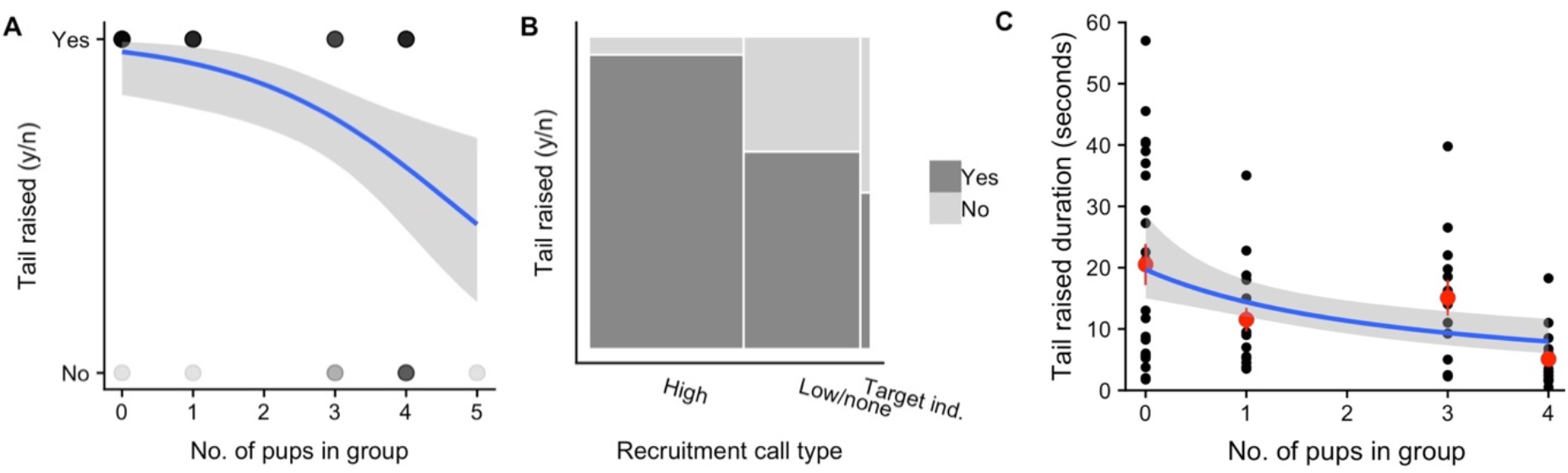
(A)The likelihood of an individual interacting with the presentation cue raising their tail yes (n = 70) or no (n = 22) dependent on the total number of pups present in the group, and (B) recruitment call type given during presentation (high urgency, low urgency or no call given, the target individual the cue was presented to). The points shading indicates the frequency of overlapping data points. Blue logistic regression line with the shaded area illustrating the 95% confidence interval. The bar surface area indicates relative frequency of response type. (C) The duration in seconds an individual raised their tail for, of the individuals that did raise their tail during an interaction with the predator cue (n = 70) dependent on the total number of pups present in the group. Red points indicate mean tail raised duration with error bars signifying standard error. Blue linear regression line with the shaded area illustrating the 95% confidence interva

#### (d) Tail raised (y/n)

Among those individuals that interacted with the predator cue, 70/92 raised their tails. GLMM analyses produced five models in the top set, of which two (model 4 and model 7: supplementary material Table 8) were retained following the application of the nesting rule. Model 4 contained only the number of pups present in the group as a negative predictor of whether an individual would raise their tail (GLMM: estimate (SE) = −0.691 (0.243), *χ*^2^= 8.418, p = 0.004; Fig.3A; supplementary material Table 2). Model 7 contained only the recruitment call type, with increased probability of individuals raising their tails following a high urgency recruitment call (estimate (SE) = 2.398 (0.818), *χ*^2^= 9.892, p = 0.007; Fig.3b; supplementary material Table 2). The number of non-pups present in the group appeared in the top set (model 13; supplementary material Table 8) having a positive effect on tail raising likelihood when included with the number of pups (GLMM; estimate (SE) = −0.691 (0.243), *χ*^2^= 0.324, p = 0.04), whereas treatment and proportion recruited (models 13, 8 and 9; supplementary material Table 8) also appeared in the top set, but did not have a robust effect (GLMM; Treatment: *χ*^2^= 7.08, p = 0.07; Proportion recruited: estimate (SE) = −1.350(1.442), *χ*^2^= 0.874, p = 0.25; supplementary material table 9).

#### (e) Tail raised duration

The duration that individuals raised their tails for ranged 0.50-57.01 seconds with a mean of 13.89±1.52 seconds. GLMM analyses produced three models in the top set, of which one (model 4; supplementary material Table 10) was retained following the application of the nesting rule. This model contained only the number of pups present in the group as a negative predictor of tail raised duration (GLMM: estimate (SE) = 0.016(0.004), *χ*^2^= 16.144, p < 0.001; Fig.3C; supplementary material Table 2). Tail raised duration was greatest when there were no pups present, 20.52±3.28 seconds, and lowest when there were the largest possible number of four pups present, 5.09±1.17 seconds. Number of non-pups and treatment (models 13 and 3; supplementary material Table 8) also appeared in the top set but did not have a robust effect (GLMM; Number of non-pups: estimate (SE) = −0.001 (0.005), *χ*^2^= 16.144, p = 0.77; Treatment: *χ*^2^= 2.22, p = 0.53; supplementary material Table 11).

#### (f) Piloerection (y/n)

Of the 92 individuals interacting with the cues, 38 individuals piloerected: 7/38 when interacting with a fur cue and 31/38 when interacting with a urine cue. GLMM analyses produced four models in the top set, of which two (model 5 and 10; supplementary material Table 12) were retained following application of the nesting rule. Model 5 contained only the predator cue type, with individuals more likely to piloerect when interacting with a urine cue than a fur cue (GLMM: estimate (SE) = 2.333(0.701), *χ*^2^= 13.542, p < 0.001; Fig.4A; supplementary material Table 2). Model 10 contained only the proportion of adults recruited as a negative predictor of whether an individual piloerected (estimate (SE) = 5.359, (1.767), *χ*^2^= 12.782, p < 0.001). Treatment did appear in the top set (model 11; supplementary material Table 12) but did not have a robust effect (*χ*^2^= 3.915, p = 0.27; supplementary material table 13). Individuals never piloerected following a low urgency or no recruitment call.

**Figure 4.**
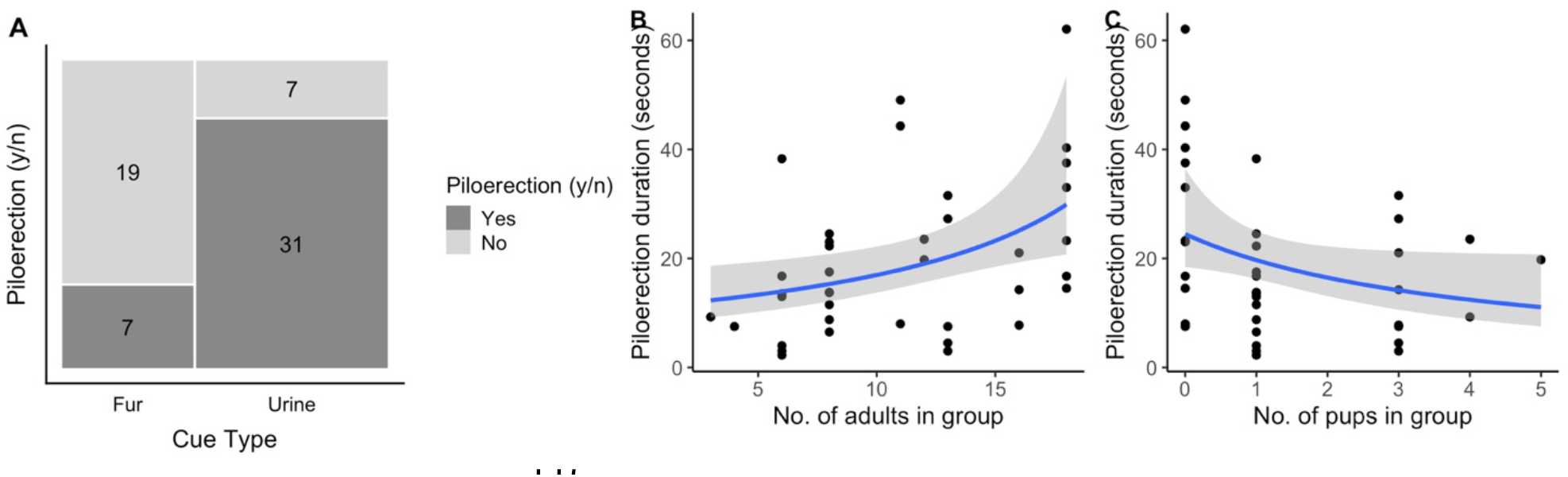
(A) The number of individuals that piloerected yes (n = 38) or no (n = 54) of those individuals interacting with the cue presentation that did piloerect for the two cue types, fur or urine. Dark grey shading indicates those individuals that did piloerect and light grey those that did not. The count for each is displayed within bar. (B) The piloerection duration for individuals interacting with the cue presentations that did piloerect (n = 38) dependent on the total number of adults present in the group and, (C) the total number of pups present in the group. Blue linear regression line with the shaded area illustrating the 95% confidence interval.

#### (g) Piloerection duration

Piloerection duration ranged from 2.25-62.01 seconds with a mean duration of 19.42±2.26 seconds. GLMM analyses produced four models in the top set, of which two (model 13 & model 2; supplementary material Table 14) were retained. Model 13 contained the number of non-pups and the number of pups present in the group. There was a positive relationship between piloerection duration and the number of non-pups (GLMM: estimate (SE) = 0.123 (0.038), *χ*^2^= 8.488, p = 0.004; Fig.4B; supplementary material Table 2). In contrast, the duration of piloerection declined as the number of pups increased (estimate (SE) = −0.189(0.060), *χ*^2^= 7.487, p = 0.006; Fig.4C; supplementary material Table 2). Model 2, containing only treatment, also appeared in the top set (*χ*^2^= 18.203, p < 0.001 supplementary material table 15). Individuals piloerected for longer durations when no pups were present (NP; 29.94±54.97 seconds; supplementary material Table 2) than in all pup present treatments: PP1 (13.45±2.67; effect size (relative to NP) = −1.17, t = −4.21, p < 0.001; supplementary material Table 11); PP2 (17.73±4.70; effect size = −0.86, t = −3.66, p < 0.001; supplementary material Table 15); PP3 (13.94±2.91; effect size = −1.13, t = −4.18, p < 0.001; supplementary material Table 15).

## Discussion

Meerkats’ mobbing-like responses towards secondary predator cues seems perplexing, given that unlike most instances of mobbing in the animal kingdom, it cannot help to drive predators away. We tested whether adults may instead use exaggerated mobbing-like responses to SPCs to teach naïve pups, but our results provided no evidence that this is the case. Contrary to our predictions, we found that adults *reduced* their mobbing-like response intensity when pups were present, particularly when more pups were present. These results strongly suggest that meerkats do not use mobbing-like responses towards SPCs as a form of teaching. Instead, we suggest that this behaviour may help to recruit other mature group members to investigate the cue and gather information to mount appropriate defensive responses.

We predicted that adults would exaggerate their mobbing-like response when pups were present and foraging with the group and that responses would be particularly exaggerated when cues were novel to pups. None of the analyses supported these predictions, as experimental treatment (NP, PP1, PP2, PP3 where PP3 was always a novel cue) did not appear to influence most of the responses investigated. There was some evidence that experimental treatment had a modest effect on interaction and piloerection duration, with interaction duration greatest when cues were novel to the group, suggestive of possible habituation through order effects. This habituation seems to have broken when a new cue (cat fur instead of cat urine, or vice versa) was presented, returning response duration to the same baseline regardless of whether pups were present. It therefore seems likely that interaction and piloerection duration were related to cue familiarity and presentation order rather than the presence or absence of pups.

Piloerection duration, an indicator of intensity, was reduced in the presence of pups irrespective of cue novelty suggesting an overall effect of pups in reducing response intensity. In the analyses of the proportion of the group recruited to inspect the SPC, whether or not interacting individuals raised their tail, and the duration of tail raising and piloerection, larger numbers of pups appeared to have an inhibitory effect on response intensity. The effect of the number of pups was reduced when the NP treatment was excluded from the analysis for the proportion of the group recruited, whether an individual raised their tail and piloerection duration, but maintained for tail raising duration (supplementary material Table 1). This suggests the presence of pups alone rather than the increasing number may drive this effect in the full data set. The reduction in response intensity could reflect the additional costs associated with provisioning pups, limiting investment in other activities. Alternatively, the high intensity of a mobbing-like response is by definition conspicuous; therefore reducing intensity when vulnerable pups are present may reduce conspicuousness and risk to pups in an area of higher perceived predation risk. Meerkats have been observed leading pups away from a predator mobbing location and therefore away from an area of increased risk (M. Manser, *pers. comm.,* February 2020). Thus, although meerkats are known to teach their pups how to hunt effectively (Thornton & McAuliffe 2006), they do not appear to use responses to SPCs to teach pups about potential predators.

If mobbing-like response to SPCs do not play a role in teaching naïve pups, what could be the function of this unusual behaviour? One possible explanation is that mobbing-like response to SPCs is a maladaptive by-product of arousal. Individuals clearly responded to the SPCs and not controls as threats, behaving similarly to how they would respond to a predator (Graw & Manser 2007). This high intensity response to SPCs may represent a misidentification of a SPC as an actual threat. However, rather than ceasing to respond to the stimuli after direct investigation, individuals tended to continue the mobbing-like behaviours whilst investigating the cues directly sniffing and scratching them. This suggests no error in classification and an awareness that the cue itself is not a threat. This cue recognition is further illustrated in the difference in response to fur versus urine cues, suggesting even a distinction within predator cue types. Interaction duration was longer for fur cues, but individuals were more likely to show the high arousal piloerection response to urine cues, possibly related to perceived risk associated. Moreover, although the mobbing-like response to SPCs is without the major costs associated with mobbing (injury, death), there are still substantial energetic, time, opportunity and conspicuousness costs of the mobbing-like response, illustrated by the reduction in response intensity potentially due to additional costs posed by pups. If there were no benefit gained from such a costly response to SPCs, it would be expected that selection would act against the persistence of this behaviour.

Arguably, a more plausible explanation is that the mobbing-like response to SPCs could play a role in information transfer. The raising of group knowledge and alertness through recruitment to SPCs can reduce risk to all members, raising vigilance and increasing speed of potential predator detection (Zöttl *et al.* 2013). A mobbing-like response may increase the probability of recruiting other group members by providing a conspicuous, localisable signal of risk. Consistent with this, our results indicate an increased probability of individuals recruiting when a higher proportion of the group is already interacting with the cue. In larger groups where individuals may be more dispersed (Focardi & Pecchioli 2005) signals may need to be more conspicuous to increase the probability of others receiving the signal. Inspection of cues may increase individual knowledge of the type of threat thus facilitating more targeted predator detection and defences. For example, stoats, *Mustela erminea*, respond with differences in scanning behaviour dependent on the source of the scent and effectiveness of the defensive response (Garvey *et al.* 2016). Previous work on meerkats has demonstrated more rapid detection of a nearby predator model following an SPC encounter (Zöttl *et al.* 2013), predator detection was not necessarily by the individual that had interacted with the cue. In addition, meerkats also show an increase in alarm calling frequency and reduce distance travelled following a natural SPC encounter (Driscoll *et al.* 2020). This supports the idea that group-level defensive responses may be enhanced by alerting conspecifics of increased risk, with recruitment further improving their knowledge of the threat.

Although meerkats do not appear to exaggerate their responses to SPCs as a form of teaching, these responses may nevertheless provide opportunities for inadvertent social learning via stimulus enhancement and/or observational conditioning. Inadvertent social learning is characterised as the transmission of learnt information between individuals without the need for experienced individuals to adjust their behaviour (Hoppitt *et al.* 2008). Meerkat pups may have sufficient inadvertent learning opportunities through observing knowledgeable group members’ high intensity responses to SPCs, without the need for exaggerated adult responses. A similar argument can be made for mobbing of actual predators: here, social learning may not be the primary adaptive function, but can be an additional benefit (Curio *et al.* 1978a; Griesser & Suzuki 2017). Whether meerkats, and other animals, learn socially from other individuals’ responses to SPCs remains to be investigated. This could be achieved by assessing whether naïve individuals’ responses towards SPCs (and the actual predators with which those SPCs are associated) change after observing a knowledgeable individual interacting with the cue.

The lack of evidence for teaching in this context may provide support for the idea that, in contrast to human teaching, which occurs across many contexts, non-human teaching is an adaptation to promote context-specific learning (Thornton & Raihani 2008). Teaching has evolved in other species when acquisition of information or a behaviour by asocial or passive social learning would be slow/dangerous or not occur at all. In the context of the mobbing-like response to SPCs, pups may have ample opportunities to learn this behaviour by watching adults’ responses, so there is no benefit for adults modifying their behaviour to promote learning. For example, meerkat pups’ responses to alarm calls become more adult-like with age, suggesting the development of experience-dependent appropriate responses to alarm calls, likely as a result of social learning, without adults altering their behaviour (Hollén & Manser 2006; Hollén *et al.* 2008). However, further research needs to be conducted on possible candidate behaviours for teaching in non-human animals to assess whether humans are the only species to perform flexible multi-context teaching.

## Acknowledgments

We thank the Kalahari Research Trust and the Northern Cape Conservation Authority for research permission (FAUNA 1020/2016). We also thank Dave Gaynor and Tim Vink for organising the field site, as well as the managers Coline Muller and Jacob Brown, and volunteers of the Kalahari Meerkat Project for organising, providing support and helping to collect the life history data and maintain habituation of the meerkats. Furthermore, we thank Michael Cant and Andrew Radford for providing valuable comments which helped improve this manuscript.

## Notes

### Competing Interest Statement

The authors have declared no competing interest.

