## Supplementary material for "Mobbing-like response to secondary predator cues is not a form of teaching in meerkats"

As the number of pups in the NP treatment was, by definition, zero, the effects of treatment and number of pups could be correlated. To address this, we also ran the analysis with the results of the NP treatment excluded. The results of these models were qualitatively very similar to those conducted on the full dataset (Table 1).

*Table 1 – Table showing the models in the top set for each aspect of response intensity, comparing the full data set with that of the dataset with the NP treatment excluded. Top models highlighted in bold following application of the nesting rule. Asterix signifies a significant effect and "NS" indicates non-significant.*

| **All conditions** | | | | **Without NP** | | | |
|---|---|---|---|---|---|---|---|
| ***(a) Proportion recruited*** | | | | | | | |
| Model | AIC | Fixed effects | Significant ? | Model | AIC | Fixed effects | Significant ? |
| **a.5** | **127.64** | **No. pups** | * | **a.4** | **94.05** | **Call type** | * |
| **a.6** | 129.26 | No. pups | * | a.6 | 95.5 | No. pups | NS |
| | | Call type | NS | | | Call type | * |
| ***(b) Interact (y/n)*** | | | | | | | |
| Model | AIC | Fixed effects | Significant ? | Model | AIC | Fixed effects | Significant ? |
| **b.10** | **236.71** | **Prop. recruited** | * | **b.10** | **176.38** | **Prop. recruited** | * |
| **b.9** | 239.06 | Prop. recruited | * | b.9 | 179.27 | Prop. recruited | * |
| | | Call type | NS | | | Call type | NS |
| **b.11** | 240.4 | Prop. recruited | * | b.11 | 179.36 | Prop. recruited | * |
| | | Treatment | NS | | | Treatment | NS |
| ***(c) Interact duration*** | | | | | | | |
| Model | AIC | Fixed effects | Significant ? | Model | AIC | Fixed effects | Significant ? |
| **c.5** | **801.22** | **Cue type** | * | c.6 | 553.3 | Treatment | * |
| **c.6** | 803.3 | Cue type | * | | | Cue type | * |
| | | Treatment | NS | **c.5** | **554.9** | **Cue type** | * |
| **c.7** | 806.1 | Treatment | * | c.3 | 558.5 | Treatment | * |
| | | No. pups | * | | | No. pups | NS |
| ***(d) Tail raised (y/n)*** | | | | | | | |
| Model | AIC | Fixed effects | Significant ? | Model | AIC | Fixed effects | Significant ? |
| **d.13** | 66.3 | No. adults | * | d.9 | 55.47 | Call type | * |
| | | No. pups | * | | | Prop. recruited | NS |

| | | | | | | | |
|---|---|---|---|---|---|---|---|
| **d.4** | **68.15** | **No. pups** | * | **d.7** | **56.38** | **Call type** | * |
| **d.7** | **68.99** | **Call type** | * | d.10 | 59.78 | Prop. recruited | NS |
| ***(e) Tail raised duration*** | | | | | | | |
| Model | AIC | Fixed effects | Significant ? | Model | AIC | Fixed effects | Significant ? |
| **e.4** | **481.73** | **No. pups** | * | **e.13** | **299.31** | **No. pups** | * |
| **e.13** | 484.04 | No. pups | * | | | **No. adults** | * |
| | | No. adults | NS | **e.5** | **300.21** | **Cue type** | * |
| **e.3** | 486.93 | No. pups | * | e.4 | 303.29 | No. pups | NS |
| | | Treatment | NS | | | | |
| ***(f) Piloerect (y/n)*** | | | | | | | |
| Model | AIC | Fixed effects | Significant ? | Model | AIC | Fixed effects | Significant ? |
| **f.5** | **93.23** | **Cue type** | * | **f.10** | **65.9** | **Prop. recruited** | * |
| **f.10** | **94** | **Prop. recruited** | * | **f.5** | **67.6** | **Cue type** | * |
| **f.11** | 96.9 | Prop. recruited | * | f.11 | 68.4 | Prop. recruited | * |
| | | Treatment | NS | | | Treatment | NS |
| ***(g) Piloerect duration*** | | | | | | | |
| Model | AIC | Fixed effects | Significant ? | Model | AIC | Fixed effects | Significant ? |
| **g.13** | **288.3** | **No. adults** | * | **g.12** | **166.6** | **No. adults** | * |
| | | **No. pups** | * | g.13 | 170.07 | No. adults | * |
| **g.2** | **292.8** | **Treatment** | * | | | No. pups | NS |
| g.12 | 292.9 | No. adults | * | | | | |

Table 1 shows that the models forming the top set were broadly consistent between the full dataset and that with the NP treatment excluded. The models in the top set for the NP-excluded dataset differ primarily only for the proportion of the group recruited, whether individuals raised their tails and piloerection duration. For these indicators of response intensity for the full dataset the number of pups had an effect, whereas when the NP treatment is excluded the numbers of pups did not have an effect. This may suggest that it is the presence of pups alone rather than the increasing number that drives this effect in the full dataset. There remains a negative effect of pup number for tail raised duration in both the full dataset and the NP-excluded data.

*Table 2. – Model summaries of the top candidate models for the indicators of mobbing intensity. (a) The proportion of the group recruited to predator cue presentation. (b) Whether an individual interacted with the cue. (c) The duration of an individual's interaction. (d) Whether an individual raised their tail. (e) The duration an individual raised their tail for. (f) Whether an individual piloerected. (g) The duration an individual piloerected for.*

| | Estimate | Std Error | Chisq | df | p value |
|---|---|---|---|---|---|
| **a. Proportion recruited** | | | | | |
| *(a.5) Number of pups* | | | | | |
| Intercept | -0.050 | 0.315 | | | |
| No. pups | -0.201 | 0.107 | 3.810 | 3 | 0.050 |
| **b. Interact (y/n)** | | | | | |
| *(b.10) Proportion of group recruited* | | | | | |
| Intercept | -1.261 | 0.269 | | | |
| Proportion recruited | 2.992 | 0.817 | 14.735 | 4 | <0.001 |
| **c. Interact duration** | | | | | |
| *(c.5) Cue type* | | | | | |
| Intercept | 3.504 | 0.139 | | | |
| Cue type | -0.511 | 0.169 | 8.787 | 5 | 0.003 |
| **d. Tail raise (y/n)** | | | | | |
| *(d.4) Number of pups* | | | | | |
| Intercept | 3.236 | 0.830 | | | |
| No. pups | -0.691 | 0.243 | 8.418 | 4 | 0.004 |
| *(d.7) Call type* | | | | | |
| Intercept | 0.547 | 0.379 | | | |
| Call type | | | 9.892 | 5 | 0.007 |
| **e. Tail raise duration** | | | | | |
| *(e.4) Number of pups* | | | | | |
| Intercept | 0.082 | 0.021 | | | |
| No. pups | 0.016 | 0.004 | 16.144 | 5 | <0.001 |
| **f. Piloerect (y/n)** | | | | | |
| *(f.5) Cue type* | | | | | |
| Intercept | -1.830 | 0.861 | | | |
| Cue type | 2.333 | 0.701 | 13.542 | 4 | <0.001 |
| *(f.10) Proportion of group recruited* | | | | | |
| Intercept | 0.611 | 1.009 | | | |
| Prop recruited | -5.359 | 1.767 | 12.782 | 4 | <0.001 |

| | | | | | |
|---|---|---|---|---|---|
| **g. Piloerect duration** | | | | | |
| *(g.13) Number of non-pups + Number of pups* | | | | | |
| Intercept | 1.571 | 0.471 | | | |
| No. non-pups | 0.123 | 0.038 | 8.488 | 6 | 0.004 |
| No. pups | -0.189 | 0.060 | 7.487 | 6 | 0.006 |
| *(g.2) Treatment* | | | | | |
| Intercept | 3.216 | 0.280 | | | |
| Treatment | | | 18.203 | 7 | < 0.001 |

### *a. Proportion of group recruited*

*Table 3. – Model selection table for the variables affecting the proportion of the adults in the group recruited to the predator cue presentation ranked by AIC value. Variables tested are the recruitment call category (high, low/none, individual presented to), treatment (NP, PP1, PP2, PP3), cue type (fur, urine), number of adults in the group, number of pups in the group. Retained models in bold.*

| Model | Fixed Effects | Intercept | Call Type | Treatment | Cue Type | No. Adults | No. Pups | df | logLik | AICc | delta | weight |
|---|---|---|---|---|---|---|---|---|---|---|---|---|
| **a.5** | **No. Pups** | **-0.050** | | | | | **-0.201** | **3** | **-60.221** | **127.64** | **0.00** | **0.463** |
| a.6 | No. Pup +Call Type | -0.226 | | | + | | -0.215 | 4 | -59.591 | 129.29 | 1.65 | 0.203 |
| a.3 | Treatment | -0.275 | | + | | | | 3 | -61.577 | 130.35 | 2.71 | 0.119 |
| a.4 | Cue Type | -0.588 | | | + | | | 3 | -61.904 | 131.01 | 3.37 | 0.086 |
| a.7 | No. Adults | -0.250 | | | | -0.020 | | 3 | -62.053 | 131.31 | 3.67 | 0.074 |
| a.2 | Call Type | -0.051 | + | | | | | 5 | -59.296 | 131.93 | 4.28 | 0.054 |
| a.1 | ALL | -0.269 | + | + | + | 0.024 | -0.275 | 9 | -57.835 | 146.53 | 18.88 | 0.000 |

### *b. Interact (y/n)*

*Table 4. – Model selection table for the variables affecting whether an individual interacts with the predator cue presentation ranked by AIC value. Variables tested are the recruitment call category (high, low, none, individual presented to), treatment (NP, PP1, PP2, PP3), cue type (fur, urine), number of adults in the group, number of pups in the group, proportion of the group interacting with the cue. Retained models in bold.*

| Model | Fixed Effects | Intercept | Call Type | Treatment | Cue Type | No. Adults | No. Pups | Prop Interact | df | logLik | AICc | delta | weight |
|---|---|---|---|---|---|---|---|---|---|---|---|---|---|
| **b.10** | **Prop Recruit** | **-1.260** | | | | | | **2.992** | **4** | **-114.245** | **236.71** | **0.00** | **0.665** |
| b.9 | Call Type + Prop Recruit | -1.233 | + | | | | | 2.695 | 6 | -113.292 | 239.06 | 2.35 | 0.206 |
| b.11 | Prop Recruit + Treatment | -0.878 | | + | | | | 2.722 | 7 | -112.879 | 240.40 | 3.68 | 0.105 |
| b.1 | ALL | -1.034 | + | + | + | 0.055 | -0.429 | 2.056 | 12 | -109.412 | 244.66 | 7.95 | 0.013 |

| | | | | | | | | | | | | |
|---|---|---|---|---|---|---|---|---|---|---|---|---|
| b.4 | No. Pups | -0.048 | | | | | -0.217 | 4 | -119.541 | 247.31 | 10.59 | 0.003 |
| b.7 | Call Type | -0.545 | + | | | | | 5 | -118.845 | 248.03 | 11.32 | 0.002 |
| b.13 | No. Adults + No. Pups | -0.427 | | | | 0.040 | -0.223 | 5 | -119.321 | 248.98 | 12.27 | 0.001 |
| b.2 | Treatment | -0.042 | | + | | | | 6 | -118.536 | 249.55 | 12.83 | 0.001 |
| b.8 | Call Type + Treatment | -0.106 | + | + | | | | 8 | -116.473 | 249.77 | 13.06 | 0.001 |
| b.3 | Treatment + No. Pups | -0.082 | | + | | | -0.245 | 7 | -117.920 | 250.48 | 13.76 | 0.001 |
| b.5 | Cue Type | -0.317 | | | + | | | 4 | -121.150 | 250.52 | 13.81 | 0.001 |
| b.6 | Cue Type + Treatment | 0.119 | | + | + | | | 7 | -118.226 | 251.09 | 14.38 | 0.001 |
| b.12 | No. Adults | -0.662 | | | | 0.017 | | 4 | -121.564 | 251.35 | 14.64 | 0.000 |

*Table 5. – Model summaries of the GLMM's appearing in the top set but not retained.*

| | Estimate | Std. Error | z value | p value |
|---|---|---|---|---|
| *b.9 Call type & Proportion recruited* | | | | |
| (Intercept) | -1.9911 | 0.7742 | 18.160 | 0.0101 |
| CallBeforehigh | 0.7584 | 0.8106 | 0.936 | 0.3494 |
| CallBeforelow | 1.0288 | 0.8358 | 1.231 | 0.2186 |
| PropRecruit | 2.6946 | 0.8446 | 3.191 | 0.0014 |
| *b.11 Treatment* | | | | |
| (Intercept) | - 0.8779 | 0.4045 | - 2.170 | 0.0210 |
| PropRecruit | 2.7220 | 0.8310 | 3.275 | 0.0011 |
| TreatmentPP1 | - 0.4050 | 0.4466 | - 0.907 | 0.3645 |
| TreatmentPP2 | - 0.7212 | 0.4717 | - 1.529 | 0.1263 |
| TreatmentPP3 | - 0.1458 | 0.4442 | - 0.328 | 0.7428 |

#### *c. Interact duration*

*Table 6. – Model selection table for the variables affecting whether an individual's interaction duration with the predator cue presentation ranked by AIC value. Variables tested are the recruitment call category (high, low, none, individual presented to), treatment (NP, PP1, PP2, PP3), cue type (fur, urine), number of adults in the group, number of pups in the group, proportion of the group interacting with the cue. Retained models in bold.*

| Model | Fixed Effects | Intercept | Call Type | Treatment | Cue Type | No. Adults | No. Pups | Prop Recruit | df | logLik | AICc | delta | weight |
|---|---|---|---|---|---|---|---|---|---|---|---|---|---|
| **c.5** | **Cue Type** | **3.504** | | | **+** | | | | **5** | **-395.263** | **801.22** | **0.00** | **0.622** |
| c.6 | Cue Type + Treatment | 3.659 | | + | + | | | | 8 | -392.781 | 803.30 | 2.07 | 0.221 |
| c.3 | Treatment + No. Pups | 3.398 | | + | | | 0.243 | | 8 | -394.207 | 806.15 | 4.93 | 0.053 |
| c.7 | Call Type | 3.496 | + | | | | | | 7 | -396.125 | 807.58 | 6.36 | 0.026 |
| c.12 | No. Adults | 3.612 | | | | -0.037 | | | 5 | -398.898 | 808.49 | 7.27 | 0.016 |
| c.2 | Treatment | 3.346 | | + | | | | | 7 | -396.785 | 808.90 | 7.68 | 0.013 |
| c.1 | ALL | 3.997 | + | + | + | -0.040 | 0.182 | -0.526 | 14 | -388.174 | 809.80 | 8.58 | 0.009 |
| c.9 | Call Type + Prop Recruit | 3.508 | + | | | | | -0.051 | 8 | -396.118 | 809.97 | 8.75 | 0.008 |
| c.10 | Prop Recruit | 3.264 | | | | | | -0.051 | 5 | -399.648 | 809.99 | 8.77 | 0.008 |
| c.4 | No. Pups | 3.245 | | | | | 0.005 | | 5 | -399.652 | 810.00 | 8.78 | 0.008 |
| c.8 | Call Type + Treatment | 3.638 | + | + | | | | | 10 | -393.862 | 810.44 | 9.22 | 0.006 |
| c.13 | No. Adults + No. Pups | 3.590 | | | | -0.037 | 0.018 | | 6 | -398.848 | 810.68 | 9.46 | 0.005 |
| c.11 | Prop Recruit + Treatment | 3.393 | | + | | | | -0.205 | 8 | -396.650 | 811.03 | 9.81 | 0.005 |

*Table 7. – Model summaries of the GLMM's containing treatment forming the top set but not retained for interaction duration.*

| | Estimate | Std. Error | t value | p value |
|---|---|---|---|---|
| *c.6 Treatment & Cue type* | | | | |

| | | | | |
|---|---|---|---|---|
| (Intercept) | 3.6587 | 0.2015 | 18.160 | 1.07E-73 |
| CueTypeUrine | -0.4993 | 0.1716 | -2.910 | 0.0036 |
| TreatmentPP1 | -0.2484 | 0.2275 | -1.092 | 0.2749 |
| TreatmentPP2 | -0.5488 | 0.2507 | -2.189 | 0.0286 |
| TreatmentPP3 | -0.1338 | 0.2140 | -0.625 | 0.5319 |
| *c.3 Treatment & Number of pups* | | | | |
| (Intercept) | 3.3981 | 0.1673 | 20.311 | 1.02E-91 |
| TreatmentPP1 | -0.7826 | 0.3508 | -2.231 | 0.0257 |
| TreatmentPP2 | -1.4774 | 0.4749 | -3.111 | 0.0019 |
| TreatmentPP3 | -0.8329 | 0.4222 | -1.973 | 0.0485 |
| NumPups | 0.2432 | 0.1065 | 2.283 | 0.0224 |

### d. *Tail raised (y/n)*

*Table 8. – Model selection table for the variables affecting whether an individual raises their tail while interacting with the predator cue presentation ranked by AIC value. Variables tested are the recruitment call category (high, low/none, individual presented to), treatment (NP, PP1, PP2, PP3), cue type (fur, urine), number of adults in the group, number of pups in the group, proportion of the group interacting with the cue. Retained models in bold.*

| Model | Fixed Effects | Intercept | Call Type | Treatment | Cue Type | No. Adults | No. Pups | Prop Recruit | df | logLik | AICc | delta | weight |
|---|---|---|---|---|---|---|---|---|---|---|---|---|---|
| d.13 | No. Adults + No. Pups | 0.777 | | | | 0.324 | -0.985 | | 5 | -27.732 | 66.37 | 0.00 | 0.456 |
| **d.4** | **No. Pups** | **3.235** | | | | | **-0.691** | | **4** | **-29.779** | **68.15** | **1.78** | **0.187** |
| **d.7** | **Call Type** | **2.944** | **+** | | | | | | **5** | **-29.042** | **68.99** | **2.62** | **0.123** |
| d.8 | Call Type + Treatment | 5.002 | + | + | | | | | 8 | -25.502 | 69.29 | 2.92 | 0.106 |
| d.9 | Call Type + Prop Recruit | 3.367 | + | | | | | -1.350 | 6 | -28.605 | 70.50 | 4.13 | 0.058 |
| d.1 | ALL | 3.742 | + | + | + | 0.779 | -0.831 | -4.179 | 12 | -21.068 | 71.42 | 5.05 | 0.036 |
| d.3 | Treatment + No. Pups | 2.996 | | + | | | -0.753 | | 7 | -29.319 | 74.39 | 8.01 | 0.008 |

| | | | | | | | | | | | |
|---|---|---|---|---|---|---|---|---|---|---|---|
| d.11 | Prop Recruit + Treatment | 5.375 | + | | | -2.915 | 7 | -29.543 | 74.84 | 8.46 | 0.007 |
| d.10 | Prop Recruit | 2.396 | | | | -1.744 | 4 | -33.239 | 75.08 | 8.70 | 0.006 |
| d.2 | Treatment | 3.569 | + | | | | 6 | -30.946 | 75.18 | 8.81 | 0.006 |
| d.12 | No. Adults | 1.069 | | | 0.072 | | 4 | -33.895 | 76.39 | 10.01 | 0.003 |
| d.5 | Cue Type | 1.847 | | + | | | 4 | -33.978 | 76.55 | 10.18 | 0.003 |
| d.6 | Cue Type + Treatment | 4.131 | + | + | | | 7 | -30.808 | 77.37 | 10.99 | 0.002 |

*Table 9. – Model summaries of the GLMM's forming the top set but not retained for tail raised (yes/no).*

| | **Estimate** | **Std. Error** | **z value** | **p value** |
|---|---|---|---|---|
| *d.8 Call type & Treatment* | | | | |
| (Intercept) | 2.2817 | 1.0488 | 2.176 | 0.0296 |
| CallPreshigh | 2.7199 | 0.9221 | 2.950 | 0.0032 |
| CallPrespres | 0.2449 | 1.5286 | 0.160 | 0.8727 |
| TreatmentPP1 | - 2.0341 | 1.3937 | - 1.459 | 0.1444 |
| TreatmentPP2 | - 2.8672 | 1.3250 | - 2.164 | 0.0305 |
| TreatmentPP3 | - 2.1859 | 1.1677 | - 1.872 | 0.0612 |
| *d.9 Call type & Proportion recruited* | | | | |
| (Intercept) | 1.0659 | 0.6832 | 1.560 | 0.1187 |
| CallPreshigh | 2.3014 | 0.8253 | 2.789 | 0.0053 |
| CallPrespres | - 0.7660 | 1.5008 | - 0.510 | 0.6098 |
| Prop. recruit | - 1.3496 | 1.4419 | - 0.936 | 0.3493 |

#### e. Tail raised duration

*Table 10. – Model selection table for the variables affecting the duration an individual's tail is raised for during an interaction with the predator cue presentation ranked by AIC value. Variables tested are the recruitment call category (high, low/none, individual presented to), treatment (NP, PP1, PP2, PP3), cue type (fur, urine), number of adults in the group, number of pups in the group, proportion of the group interacting with the cue. Retained models in bold.*

| Model | Fixed Effects | Intercept | Call Type | Treatment | Cue Type | No. Adults | No. Pups | Prop Recruit | df | logLik | AICc | delta | weight |
|---|---|---|---|---|---|---|---|---|---|---|---|---|---|
| **e.4** | **No. Pups** | **0.082** | | | | | **0.016** | | **5** | **-235.395** | **481.73** | **0.00** | **0.668** |
| e.13 | No. Adults + No. Pups | 0.096 | | | | -0.001 | 0.015 | | 6 | -235.352 | 484.04 | 2.31 | 0.210 |
| e.3 | Treatment + No. Pups | 0.090 | | + | | | 0.032 | | 8 | -234.285 | 486.93 | 5.20 | 0.050 |
| e.6 | Cue Type + Treatment | 0.102 | | + | + | | | | 8 | -235.457 | 489.27 | 7.55 | 0.015 |
| e.8 | Call Type + Treatment | 0.057 | + | + | | | | | 9 | -234.170 | 489.34 | 7.61 | 0.015 |
| e.5 | Cue Type | 0.132 | | | + | | | | 5 | -239.477 | 489.89 | 8.16 | 0.011 |
| e.2 | Treatment | 0.078 | | + | | | | | 7 | -237.067 | 489.94 | 8.21 | 0.011 |
| e.12 | No. Adults | 0.170 | | | | -0.007 | | | 5 | -239.685 | 490.31 | 8.58 | 0.009 |
| e.11 | Prop Recruit + Treatment | 0.071 | | + | | | | 0.027 | 8 | -236.702 | 491.76 | 10.04 | 0.004 |
| e.1 | ALL | 0.142 | + | + | + | -0.006 | 0.028 | 0.052 | 13 | -229.655 | 491.81 | 10.08 | 0.004 |
| e.7 | Call Type | 0.080 | + | | | | | | 6 | -240.654 | 494.64 | 12.91 | 0.001 |
| e.9 | Call Type + Prop Recruit | 0.072 | + | | | | | 0.028 | 7 | -240.472 | 496.75 | 15.02 | 0.000 |
| e.10 | Prop Recruit | 0.095 | | | | | | 0.017 | 5 | -243.365 | 497.67 | 15.94 | 0.000 |

*Table 11. – Model summaries of the GLMM's forming the top but not retained for tail raised duration.*

| | Estimate | Std. Error | t value | p value |
|---|---|---|---|---|

| e.3 Treatment & Number of pups | | | | |
|---|---|---|---|---|
| (Intercept) | 0.0897 | 0.0230 | 3.899 | 9.64e-05 |
| TreatmentPP1 | - 0.0359 | 0.0408 | - 0.882 | 0.3780 |
| TreatmentPP2 | - 0.0643 | 0.0460 | - 1.398 | 0.1621 |
| TreatmentPP3 | - 0.0432 | 0.0435 | - 0.992 | 0.3211 |
| NumPups | 0.0318 | 0.0141 | 2.256 | 0.0241 |

### f. Piloerection (y/n)

*Table 12. – Model selection table for the variables affecting whether an individual piloerects during an interaction with the predator cue presentation ranked by AIC value. Variables tested are the treatment (NP, PP1, PP2, PP3), cue type (fur, urine), number of adults in the group, number of pups in the group, proportion of the group interacting with the cue. Models containing recruitment call category were not included because of convergence issues due to no individuals piloerecting for the low/none category. Retained models in bold.*

| Model | Fixed Effects | Intercept | Treatment | Cue Type | No. Ad | No. Pups | Prop Recruit | df | logLik | AICc | delta | weight |
|---|---|---|---|---|---|---|---|---|---|---|---|---|
| f.1 | ALL | -1.669 | + | + | 0.147 | -1.991 | -9.050 | 10 | -28.695 | 80.11 | 0.00 | 0.997 |
| **f.5** | **Cue Type** | **-1.830** | | **+** | | | | **4** | **-42.387** | **93.23** | **13.13** | **0.001** |
| **f.10** | **Prop Recruit** | **0.611** | | | | | **-5.359** | **4** | **-42.768** | **94.00** | **13.89** | **0.001** |
| f.11 | Prop Recruit + Treatment | 1.157 | + | | | | -5.949 | 7 | -40.810 | 96.95 | 16.85 | 0.000 |
| f.6 | Cue Type + Treatment | -1.721 | + | + | | | | 7 | -41.945 | 99.22 | 19.12 | 0.000 |
| f.4 | NumPups | 0.054 | | | | -0.323 | | 4 | -47.695 | 103.85 | 23.74 | 0.000 |
| f.13 | No. Ad + No. Pups | -1.227 | | | 0.144 | -0.327 | | 5 | -47.181 | 105.06 | 24.95 | 0.000 |
| f.3 | Treatment + No, Pups | -0.335 | + | | | -0.970 | | 7 | -45.130 | 105.59 | 25.49 | 0.000 |
| f.12 | No. Ad | -1.953 | | | 0.157 | | | 4 | -48.573 | 105.61 | 25.50 | 0.000 |
| f.2 | Condition | -0.168 | + | | | | | 6 | -47.592 | 108.17 | 28.07 | 0.000 |

*Table 13. – Model summaries of the GLMM's forming the top but not retained for piloerection (yes/no).*

| | Estimate | Std. Error | z value | p value |
|---|---|---|---|---|
| *f.11 Proportion recruited & Treatment* | | | | |
| (Intercept) | 1.1572 | 1.1633 | 0.995 | 0.3199 |
| Prop. recruit | - 5.9486 | 1.9152 | - 3.106 | 0.0019 |
| TreatmentPP1 | 0.0221 | 0.8673 | 0.026 | 0.9796 |
| TreatmentPP2 | - 0.4800 | 0.9021 | - 0.532 | 0.5946 |
| TreatmentPP3 | - 1.4624 | 0.8773 | - 1.667 | 0.0955 |

### g. *Piloerection duration*

*Table 14. – Model selection table for the variables affecting the duration an individual piloerected for during an interaction with the predator cue presentation ranked by AIC value. Variables tested are the recruitment call category (high, low/none, individual presented to), treatment (NP, PP1, PP2, PP3), cue type (fur, urine), number of adults in the group, number of pups in the group, proportion of the group interacting with the cue. Retained models in bold.*

| Model | Fixed Effects | Intercept | Call Type | Treatment | Cue Type | No. Adults | No. Pups | Prop Recruit | df | logLik | AICc | delta | weight |
|---|---|---|---|---|---|---|---|---|---|---|---|---|---|
| **g.13** | **No. Adults + No. Pups** | **1.571** | | | | **0.123** | **-0.189** | | **6** | **-136.784** | **288.28** | **0.00** | **0.731** |
| **g.2** | **Treatment** | **3.216** | | **+** | | | | | **7** | **-137.569** | **292.87** | **4.59** | **0.074** |
| g.12 | No. Adults | 0.582 | | | | 0.206 | | | 5 | -140.528 | 292.93 | 4.65 | 0.071 |
| g.4 | No. Pups | 2.906 | | | | | -0.251 | | 5 | -141.028 | 293.93 | 5.65 | 0.043 |
| g.6 | Cue Type + Treatment | 3.003 | | + | + | | | | 8 | -137.064 | 295.09 | 6.81 | 0.024 |
| g.11 | Prop Recruit + Treatment | 3.383 | | + | | | | -0.762 | 8 | -137.208 | 295.38 | 7.10 | 0.021 |
| g.8 | Call Type + Treatment | 3.298 | + | + | | | | | 8 | -137.269 | 295.50 | 7.22 | 0.020 |
| g.3 | Condition + No. Pups | 3.241 | | + | | | 0.037 | | 8 | -137.532 | 296.03 | 7.75 | 0.015 |

| | | | | | | | | | | | | | |
|---|---|---|---|---|---|---|---|---|---|---|---|---|---|
| g.10 | Prop Recruit | 2.912 | | | | | | -1.122 | 5 | -145.709 | 303.29 | 15.01 | 0.000 |
| g.9 | Call Type + Prop Recruit | 3.450 | + | | | | | -2.365 | 6 | -144.751 | 304.21 | 15.93 | 0.000 |
| g.1 | ALL | 2.368 | + | + | + | 0.082 | -0.060 | -1.375 | 12 | -134.164 | 304.81 | 16.53 | 0.000 |
| g.5 | Cue Type | 2.539 | | | + | | | | 5 | -146.553 | 304.98 | 16.70 | 0.000 |
| g.7 | Call Type | 2.648 | + | | | | | | 5 | -146.653 | 305.18 | 16.90 | 0.000 |

*Table 15. – Model summaries of the GLMM's containing treatment forming the top set for piloerection duration.*

| | **Estimate** | **Std. Error** | **t value** | **P value** |
|---|---|---|---|---|
| *g.2 Treatment* | | | | |
| (Intercept) | 3.2161 | 0.2800 | 11.487 | < 0.001 |
| TreatmentPP1 | -0.8084 | 0.1922 | -4.206 | < 0.001 |
| TreatmentPP2 | -1.1039 | 0.3016 | -3.660 | < 0.001 |
| TreatmentPP3 | -1.1123 | 0.2663 | -4.177 | < 0.001 |
